# Metatranscriptomics and nitrogen fixation from the rhizoplane of maize plantlets inoculated with a group of PGPRs

**DOI:** 10.1101/437087

**Authors:** Lorena Jacqueline Gómez-Godínez, Ernesto Ormeño-Orrillo, Esperanza Martínez-Romero

## Abstract

The free-living soil bacteria that are beneficial for the growth of plants are known as plant growth-promoting rhizobacteria (PGPR). In this work, a multi-species of PGPR bacteria inoculant was designed, which included nitrogen-fixing strains such as *Rhizobium phaseoli*, *Sinorhizobium americanum* and *Azospirillum brasilense*, as well as other plant growth promoting bacteria such as *Bacillus subtillis* and *Methylobacterium extorquens*. The multi-species community exerted a beneficial effect on plant seedlings when it was inoculated, greater than the effect observed when inoculating each bacteria individually. Acetylene reduction of maize roots was recorded with the multi-species inoculant, which suggests that nitrogen fixation occurred under these conditions. To analyze the contributions of the different nitrogen-fixing bacteria that were inoculated, a metatranscriptomic analysis was performed. The differential expression analysis revealed that the predominantly *nif* transcripts of *Azospirillum* are overexpressed, suggesting that it was responsible for nitrogen fixation in maize. Overall, we analyzed the interaction of a synthetic community, suggesting it as an option, for future formulations of biofertilizers.

**IMPORTANCE:** While nodulation processes and nitrogen fixation by rhizobia have been well studied, little is known about the interaction between rhizobia and non-leguminous plants such as maize, which is used as a model for this study. Nitrogen fixation in cereals is a long searched goal. Instead of single species inoculants, multi-species inoculation may be more efficient to promote plant growth and fix nitrogen. Metatrascriptomes allowed us to recognize the bacteria responsible for nitrogen fixation in plant rootlets. The study of the function of certain genes may help to understand how microorganisms interact with the root plant, as well as allow a better use of microorganisms for the generation of novel biofertilizers using microbial consortia.

## INTRODUCTION

In nature and in agricultural fields there is a large diversity of bacteria associated with plants. Once isolated in culture media and subsequently tested individually in plant assays, many plant-bacteria prove to be capable of promoting plant growth. Bacterial mechanisms of action are diverse, and some bacteria may exhibit more than one mechanism. Plant growth promoting rhizobacteria (PGPR) may produce phytohormones or volatiles, solubilize nutrients, fix nitrogen or inhibit pests/pathogens. Among them, nitrogen-fixing bacteria are in general a minority, possibly due to the metabolic charge of fixing nitrogen.

As bacteria are not alone in soil and in plants, recent inoculation assays have considered the use of more than single strains, and combinations of *Azospirillum* and rhizobia or *Bacillus* and rhizobia have been more successful than single-strain inoculants in promoting plant growth (reviewd in 1–3). For example, *Azospirillum* and *Bradyrhizobium* inoculation enhances nodulation and nitrogen fixation in legumes such as soybean (4) and, in chickpea a combined inoculation of *Mesorhizobium*, *Pseudomonas* and the *Piriformospora indica* fungi increased 31% plant dry weight. *Azospirillum* coinoculation with rhizobacteria may alleviate plant stress (5).

### Multispecies inoculum bacteria

#### Azospirillum

Among the PGPRs, *Azospirillum* deserves special attention. It was one of the first diazotrophs isolated from non-legumes (6, 7) but its role for fixing nitrogen was critically dismissed. A revision of plant growth promotion mechanisms in *Azospirillum* has been reported (8). A global analysis of 30 years of *Azospirillum* trials in agricultural crops including maize showed and average 10-30% increase in yield when inoculated (9). In a controlled assay under hydric stress, *Azospirillum* in maize plants increased 10% the number of sprouts and 7% of plant height (10). *Azospirillum* is widely used as a commercial product. Auxin production by *Azospirillum* is considered the main mechanism to increase plant growth (11, 12), and three different enzymatic pathways to produce auxins have been described in *Azospirillum* (7), mutants that overproduce auxins have enhanced effects (13).

#### Bacillus

*Bacillus* strains may produce auxins (14) as well as many antibiotics, and has been reported to enhance plant growth (15–17). A large diversity of *Bacillus* were found in roots of *Phaseolus vulgaris*, derived from seeds (18). As *P. vulgaris* and maize are grown in associated crops (19), we surmise that they share their *Bacillus* symbionts and other root. *Methylobacterium* are found in many plants in roots, leaves and stems (20), and are capable to produce cytokinins and metabolize methanol that is produced as a side product during plant wall synthesis. It promotes growth in plants and mosses (21).

### Rhizobia

Rhizobia are the best studied plant symbionts which have been used in agriculture for over 120 years. Rhizobia are used in commercial inoculants for different crops, especially soybean. Rhizobia form nodules and fix nitrogen. In nodules, rhizobia may fix over 250 kg of N per ha per year equivalent to a heavy chemical fertilization of crops. Additionally, rhizobia may colonize the rhizospheres of many plants and the inside tissues of non-legumes. Rhizobia may exert beneficial effects in non-legume plants similar to PGPRs. In *Sinorhizobium*, *nif* genes that are needed for free living nitrogen fixation were found (22) and thus we considered it as a good candidate for nitrogen fixation in non-legumes.

Nitrogen fixation in cereals has been long pursued since cereals such as rice, maize and wheat became widely consumed worldwide (23), considering they use the largest amounts of chemical nitrogen fertilizers which represent a substantial cost for farmers, not to mention the high toll these fertilizers take on the environment. Maize originated and was domesticated in Mexico where many native races and varieties exist (24). Here we used a native Mexican race for the multispecies inoculation experiment.

A differential gene regulation of bacteria occurs in plants and has been reported in *Bacillus* (25), *Azospirillum* (26), *Sinorhizobium* (27), and *Rhizobium* (28) in single bacterium inoculation assays. *Bacillus* in maize exudates has increased expression of genes for nutrient utilization, synthesis of antimicrobial peptides and of chemotaxis and motility (29). *Azospirillum brasilense* showed upregulation of genes involved in biofilm formation, chemotaxis and nitrogen fixation in roots of wheat (26). In rice roots, *Azospirillum lipoferum* expressed genes related ROS detoxification and multidrug efflux pumps but *nif* genes were not induced (30). An antioxidant response has also been described in *Methylobacterium* when exposed to soybean root exudates (31). In rhizobia genes expressed in non-legume roots are involved in the transport of root derived nutrients and in legumes defense to plant phytoalexins (32). Furthermore, bacterial gene expression may be altered in presence of other root bacteria. Here we analyzed the metatranscriptome of a synthetic bacterial community in the roots of a native maize landrace. Considering that there were three nitrogen fixing bacteria in the inoculant it was of interest to determine which, if any would fix nitrogen in maize.

## MATERIALS AND METHODS

### Bacterial strains and cultures

*Rhizobium phaseoli* Ch24-10 (33), *Azospirillum brasilense* sp.7 (6), *Bacillus subtilis* CCGE2031 (18), *Methylobacterium extorquens* am1 (34) and *Sinorhizobim americanum CFNEI156* (35), were grown in PY medium at 30°C for 12 h. Cells were harvested by centrifugation at 6000 rpm for 15 min, resuspended in MgSO_4_ 10mM and adjusted to a final concentration of 10^8^ cfu ml^−1^.

### Inhibition of growth between multi-species inoculum

To verify that the chosen community could grow together and there was no inhibition of growth among them we did antibiosis tests. Individual strains were grown in liquid PY medium at 30°C for 12 h. Cells were harvested by centrifugation at 6000 rpm for 15 min, resuspended in MgSO_4_ 10mM and adjusted to a final concentration of 10^8^ cfu ml^−1^. Harvested cells from each strain were then dripped on solid PY medium at 30°C and monitored for 8 days. The treatment was done by biological triplicate and a qualitative result was obtained.

### Cultivation in PY medium of the multi-species inoculum for transcriptome

Cultures were made by inoculating 30 ml of PY medium with *Rhizobium phaseoli* Ch24-10, *Azospirillum brasilense* sp7, *Methylobacterium extorquens* AM1, *Bacillus subtilis* CCGE2031 and *Sinorhizobium americanum* CFNEI 156, as well as *R. phaseoli* Ch24-10 alone were inoculated only at a final bacterial concentration of 10^8^, treatments were performed in triplicate. After 24 hours, 3 ml of RNAlater (Ambion) were added, the samples were centrifuged for 15 minutes at 6000 rpm at 4 ° C, a pellet was obtained from the cultures and RNA was extracted (Fig. 1).

**Figure 1.**
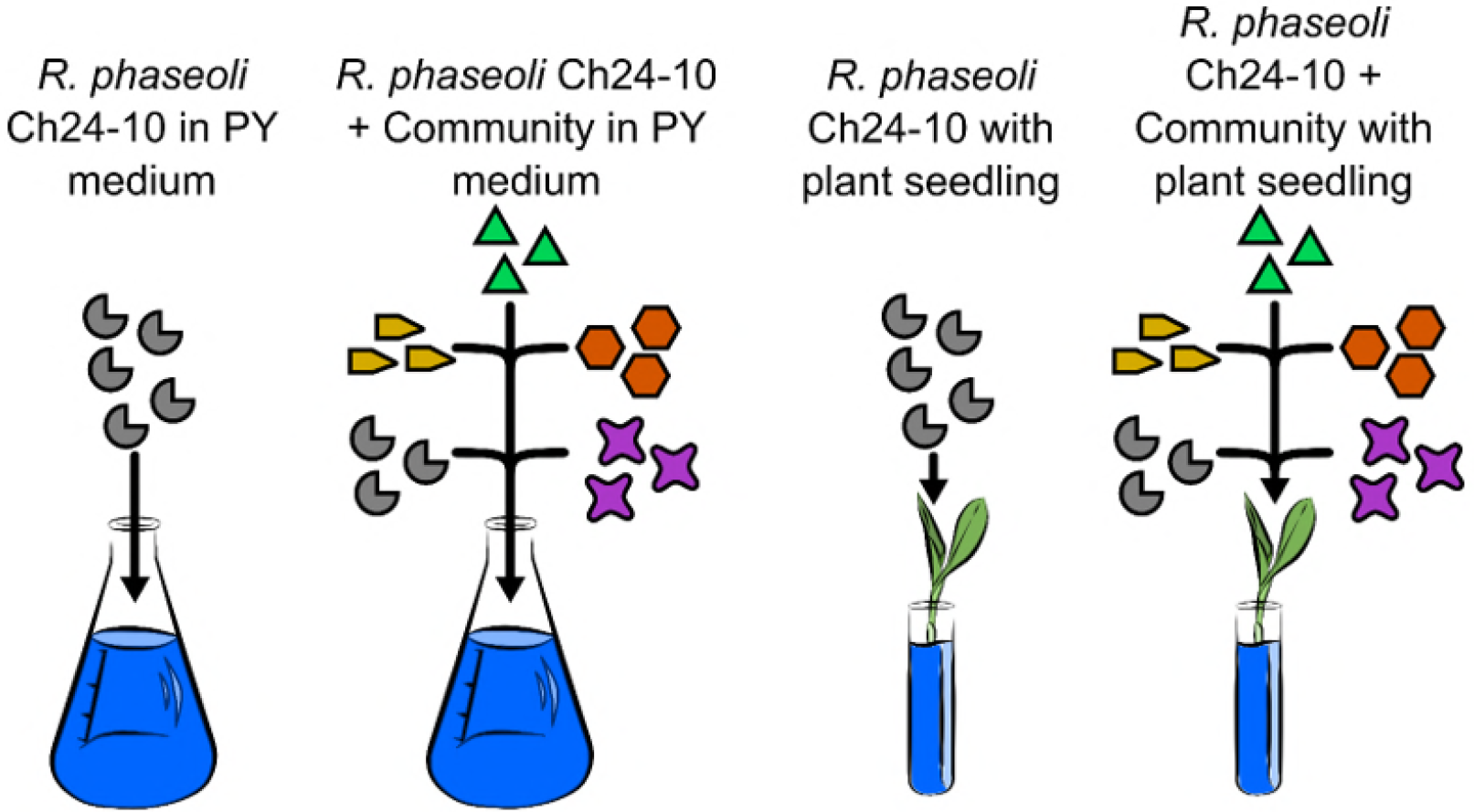
Overview of the workflow to determine changes within the transcriptomes of *Rhizobium phaseoli* Ch24-10 grown PY medium and plant with and without a bacterial community. After 24 hours, for cultures in PY medium and 5 days in cultures made with plants after inoculation, RNA extraction was carried out, extracted RNA was sequenced and the analysis of the data was performed as described in the Methods section.

From the pellet obtained from the PY medium, RNA extraction was performed through the Qiagen protocol. The RNA obtained was quantified by Nanodrop 2000 (Thermo Scientific). The integrity of the total RNA was verified by means of 1% agarose gels and in Bioanalyzer 2100 (Agilent Technologies).

### Germination, inoculation and growth of seedlings

Black Criollo maize seeds were surface disinfected with 70% ethanol (1 min), 30% sodium hypochlorite (20 min), 2% thiosulfate (2 min) and rinsed thoroughly with sterile water. Disinfected seeds were germinated in agar-water plates in the dark for 24 h at 30°C, and subsequently transplanted into flasks with sterile vermiculite. Plants were irrigated with nitrogen free Fahraeus solution (0.132 g/L CaCl_2_, 0.12 g/L MgSO_4_.7H_2_O, 0.1 g/L KH_2_PO_4_, 0.075 g/L Na_2_HPO_4_.2H_2_O, 5 mg/L Fe-citrate, and 0.07 mg/L each of MnCl_2_.4H_2_O, CuSO_4_.5H_2_O, ZnCl_2_, H_3_BO_3_, and Na2MoO4.2H2O, adjusted to pH 7.5 before autoclaving (36), and inoculated with 10^8^ cfu ml^−1^ of each bacteria; single inoculants and plants without inoculants were used as controls. Plants were maintained for 30 days at 28 °C with a 12 h dark/light period. At 30 days after inoculation, growth of the aerial part, fresh and dry weight and chlorophyll content were measured.

For RNA extraction, *in vitro* plant cultures were carried out under hydroponic and axenic conditions. Seedlings from disinfected seeds were transferred to glass tubes (one seedling in each tube) containing 10 mL of Fahraeus medium. Plants were inoculated with 10^8^ bacteria, and single inoculants and plants without inoculants were used as controls. Plants were maintained for 5 days at 28°C with a 12 h dark light/period. The data obtained from each of the treatments were analyzed with an analysis of variance (ANOVA) and the comparison test of means Tukey (p = 0.05) (Fig. 1).

### RNA extraction, RNA-seq library construction and sequencing

Root-attached bacteria were recovered from hydroponic cultures 5 days after inoculation by sonicating roots for 15 min from 50 seedlings in 5 mL of RNAlater stabilization solution (Ambion), followed by centrifugation (6000 rpm; 15 min; 4°C). The cell pellet was resuspended in RNAlater solution. Total RNA was extracted using a Qiagen kit and treated with DNAse (Qiagen) following the manufacturer instructions.

Sequencing was done at the Genomics and Bioinformatics Service Texas A&M AgriLife. rRNA was depleted using the Ribo-Zero Bacteria Protocol (Illumina). Sequencing libraries were prepared using the protocol TruSeq Stranded (Illumina) and sequenced with Illumina HiSeq 2500 (2×125).

Lastly, twelve libraries were obtained three from rhizobium alone and three from rhizobium with the community in PY and three from rhizobium alone and three from rhizobium with the community in plant. The sequencing data are available in the Texas A&M AgriLife website. (https://download.txgen.tamu.edu/Mateos/150804_D00572_0126_AC79VYANXX_15035Mts/).

### Data analyses

Raw sequencing data were scanned for adapters and quality assessed using FASTQC (v0.11.2) (37). Sequencing adapters and low quality sequences were removed using Trimmomatic (v0.38) (38) with the parameters ILLUMINACLIP:Adapter.fasta:2:30:7 SLIDINGWINDOW:4:15 MINLEN:60. Paired-end sequencing reads were then mapped to concatenated genome sequence that contains all the members of the community using Bowtie2 (v2.1.0) (39), with the following parameters --sensitive --rdg 1000,1000 --rfg 1000,1000 1 -k 20. Gene abundance was quantified using featureCounts from the Subread software (v1.6.2) (40) using the following parameters -T 8 -p -a -t CDS -g ID -o./. Genes that did not have at least 1 cpm in at least one sample were removed from further analysis (Supplementary 2).

The R package edgeR (v3.22.3)(41), was used to test for differential gene expression, considering a gene as differentially expressed at FDR < 0.05. Functional annotation was performed using Trinotate (v3.0.1), and GO term enrichment was performed using the topGO R package (v2.32.0) (42), using a Fisher classic test and setting an enrichment threshold of p-value < 0.05 (Supplementary 2).

## RESULTS

### The multi-species inoculum improved growth of maize

Before designing the multispecies inoculum, we evaluated whether the microbial consortium was able to grow together and was cultivated in solid PY medium, observing that there was no inhibition between any of the chosen bacteria (data no shown).

A model of plant-bacterial interaction was set up under axenic conditions. Maize seedlings were incubated at 28°C for 5 days with a light cycle of 12 hours and then inoculated with bacterial suspension containing 10^8^ of the inoculum multi-species, other treatment with only strain as inoculum and controls were non-inoculated maize seedlings.

Shoot weight was higher in plants inoculated with the multispecies group in comparison to non-inoculated plants or plants inoculated with single bacterial species (Fig. 2), with increases up to over 100%. Chlorophyll content increased 60%, as well as in plants with the multispecies inoculum in comparison to non-inoculated plants or plants inoculated with single bacterial species (Fig. 2).

**Figure 2.**
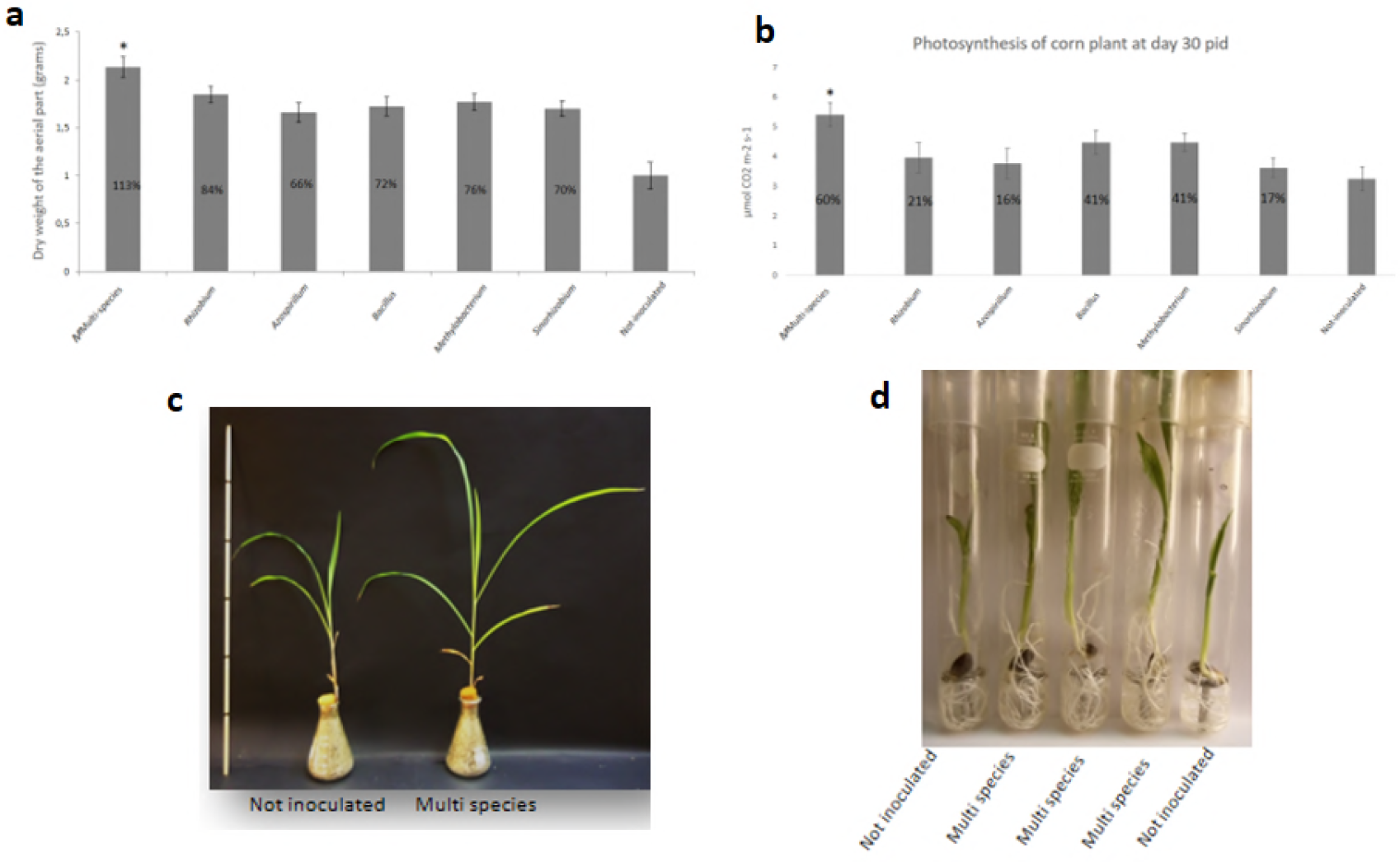
Effect of multi-species inoculum on maize growth (a) Aerial mass of the plant thirty days after inoculation (b) Improvement in photosynthetic rate in maize plants inoculated 30 days after colonization (c) Comparison of an uninoculated plant and a plant colonized with the multi-species inoculum thirty days after inoculation (d) Increase in root mass in plants colonized with the multi-species inoculum. The numbers inside the bars represent the percentage of gain compared to the control statistically significant Tukey (value of p <0.05).

### RNA-seq profiling of *Rhizobium phaseoli* Ch24-10 interacting with maize seedlings

We used the FastQC tool to visualize the quality of the sequences, and in general the samples were of good quality. The RNA-Seq generated approximately 40 million reads on average for each sample, of which 98% were confirmed to be valid after filtering reads using Trimmomatic (supplementary 1).

For gene differential expression analyses, reads were first mapped to the *R. phaseoli* genome or the concatenated sequence of the genomes of the members of the community using Bowtie2, with the featureCounts software, we counted the reads mapped in the different contigs and obtained the matrix of accounts that was used for the differential expression. The Spearman correlation coefficient of the transcriptome data for the inoculated libraries of *R. phaseoli* alone and *R. phaseoli* with community was 0.95 on average, indicating that the biological and technical replicas had a high reproducibility.

### Genes involved with nitrogen fixation are activated when *Rhizobium* interacts with the corn plant

Our analyses revealed 730 up-regulated (fold-change higher than 1 and FDR ≤ 0.05) and 376 down-regulated genes when comparing samples of *R. phaseoli* Ch24-10 grown in PY medium against *R. phaseoli* Ch24-10 grown in plants. Within the up-regulated genes, there are genes involved with transport, and encode ABC transporter permease, urea ABC transporter ATP-binding protein UrtD and cation transporter.

Genes involved with cellular respiration were also found as up-regulated in *R. phaseoli.* They encode T cbb3-type cytochrome c oxidase subunit FixP, cbb3-type cytochrome oxidase assembly protein CcoS, cytochrome b, cytochrome c, cytochrome o ubiquinol oxidase subunit I, cytochrome o ubiquinol oxidase subunit IV, cytochrome-c oxidase, cbb3-type subunit II, and cytochrome P450.

Many of the genes that were up-regulated in this condition are associated with nitrogen fixation. They encode, for example, nitrogen fixation protein NifX, nitrogen fixation protein NifZ, nitrogenase, nitrogenase cofactor biosynthesis protein NifB, nitrogenase iron-molybdenum cofactor biosynthesis protein NifN, nitrogenase stabilizing/protective protein NifW were found to be upregulated when *R. phaseoli* was grown in seedlings (Supplementary 3).

Within the genes that were found down-regulated were genes involved with chemotaxis and encode chemotaxis proteins CheR and CheW as well as chemotaxis response regulator protein-glutamate methylesterase; and genes involved in flagellar synthesis processes, such as those encoding flagellar basal body protein FliL, flagellar basal body rod protein FlgC, flagellar basal body rod protein FlgG, flagellar hook-basal body complex protein FliE, flagellar motor switch protein FliN, flagellin, flagellin C protein, flagellin synthesis regulator protein and flagellum biosynthesis repressor protein FlbT (Supplementary 3).

When comparing *R. phaseoli* Ch24-10 in plant against *R. phaseoli* Ch24-10 that was inoculated with the bacterial community in plant seedlings, we found 110 up-regulated and 53 down-regulated genes. Encompassed among up regulated genes were genes involved with transport, encoding ABC transporter ATP-binding protein, ABC transporter permease, ABC transporter substrate-binding protein, MFS transporter, peptide ABC transporter, sugar ABC transporter ATP-binding protein, and many hypothetical genes (Supplementary 3).

Among the downregulated genes were genes involved with respiration such as those encoding cytochrome P450 and electron transfer flavoprotein subunit alpha/FixB family protein, as well as fixing nitrogen proteins such as nitrogen fixation protein, nitrogen fixation protein NifZ, nitrogenase cofactor biosynthesis protein NifB and nitrogenase iron-molybdenum cofactor biosynthesis protein NifN (Supplementary 4).

Notably when *Rhizobium* is the only bacterial inoculant its machinery for nitrogen fixation is expressed. However, when *Rhizobium* is inoculated along the community, this machinery is repressed, and the role is overtaken by other members of the community. The results imply that *Azospirillum* overtakes the nitrogen fixation function, observed mainly in the overexpression of *Azospirillum* genes involved in said function in both PY medium or in plants (Fig. 3).

**Figure 3.**
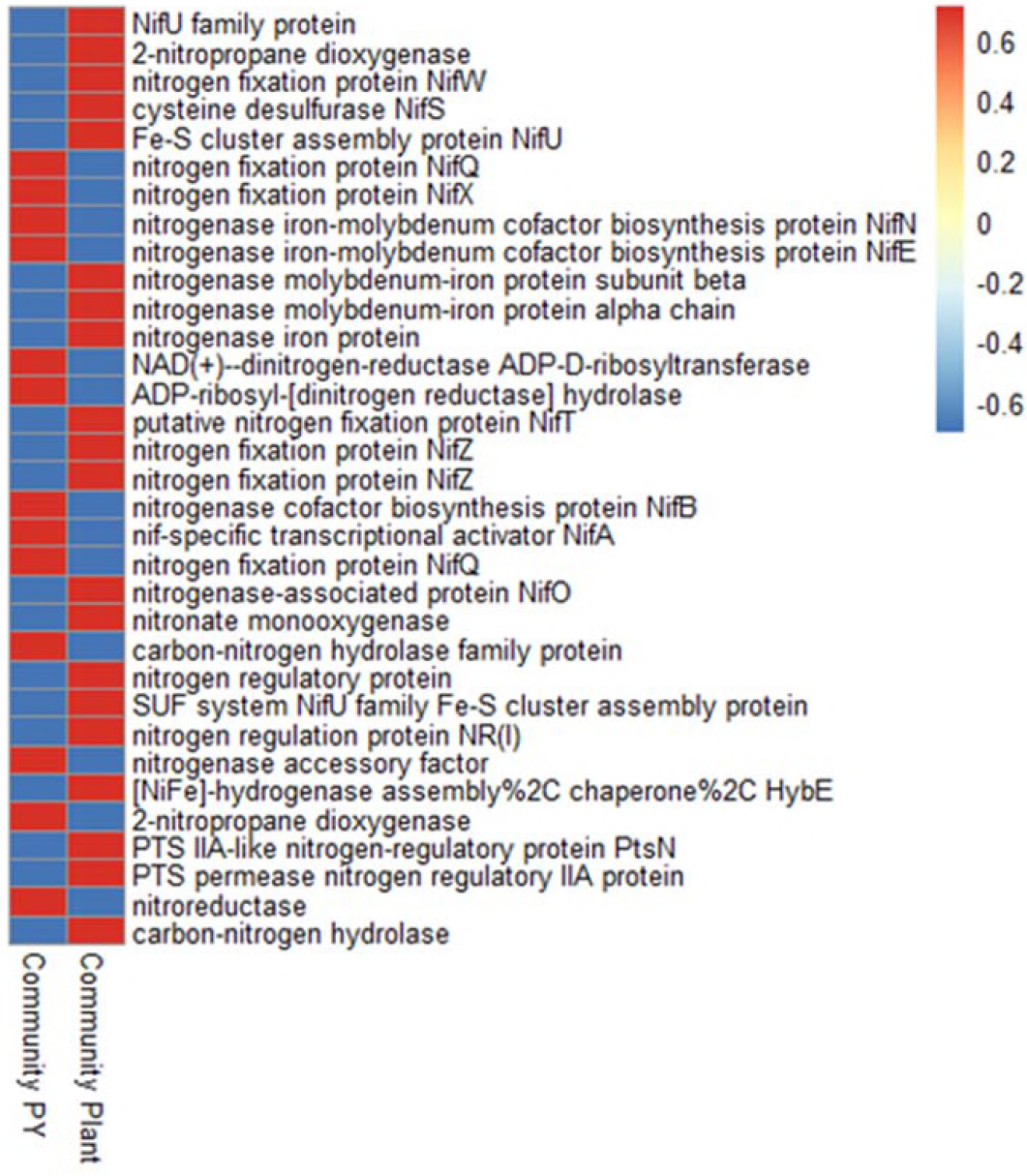
Heatmap of global gene expression profiles of *nif* genes in *Azospirillum brasilense* sp7. Expression values were represented as a Z-score due to the difference of expression among genes.

This differentially expressed genes enriched with the topGO tool and represented with REViGO are shown in the Figure 4 and 5.

**Figure 4.**
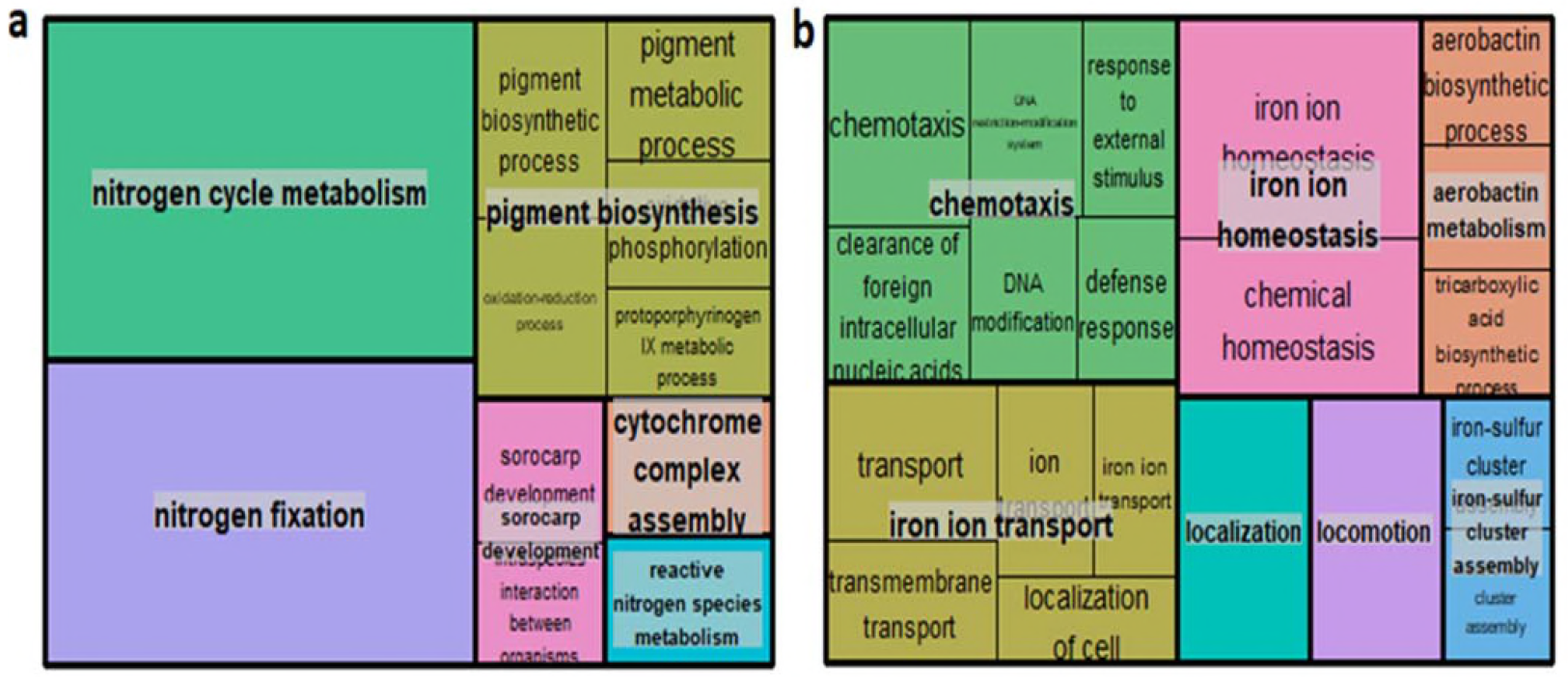
Visualization of GO terms. Treemaps of enriched Biological Process GO terms obtained using REVIGO when comparing *R. phaseoli* grown in PY medium against *R. phaseoli* Ch24-10 grown in plants, using a) up-regulated genes b) down-regulated genes.

**Figure 5.**
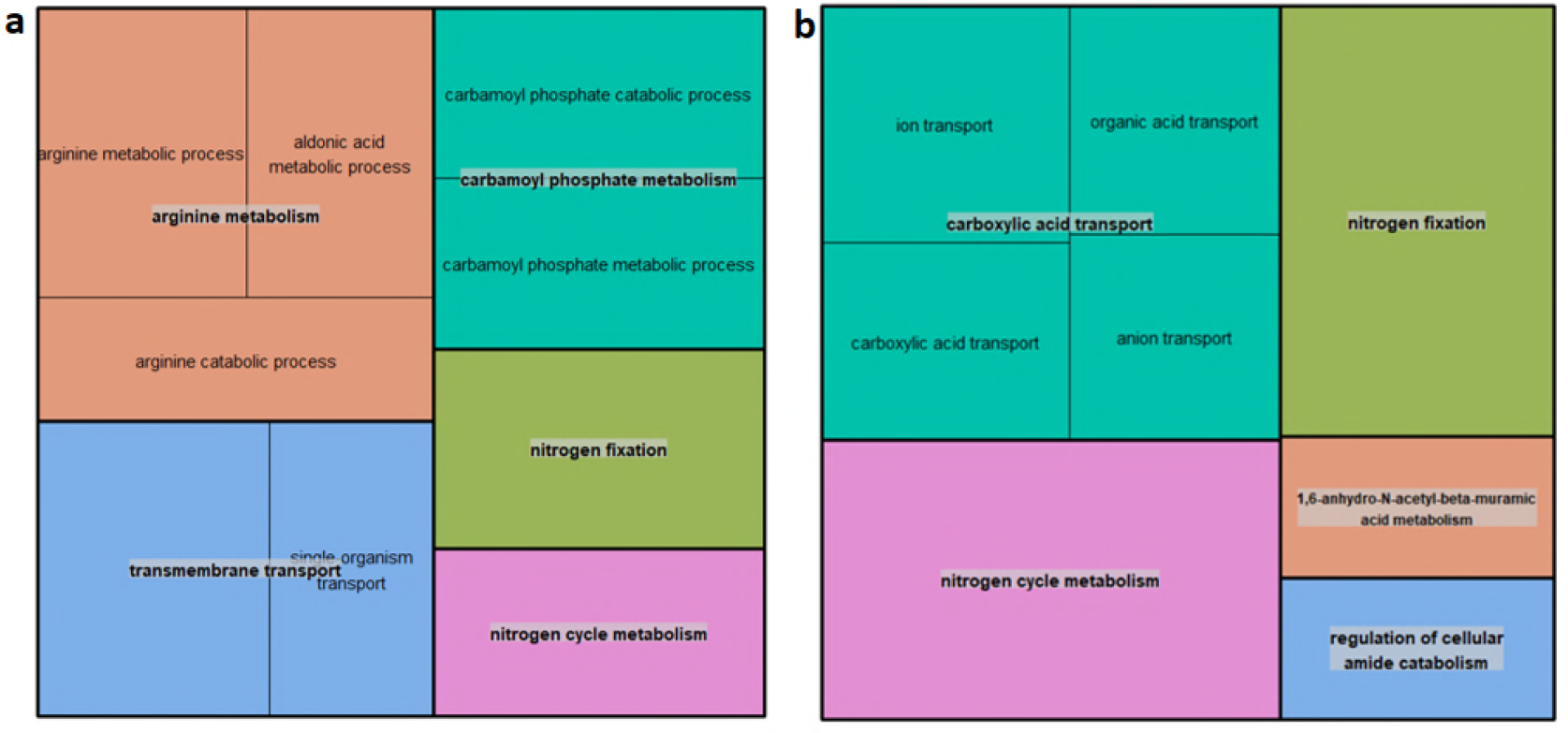
Visualization of GO terms. Treemaps of enriched Biological Process GO terms obtained using REVIGO when comparing *R. phaseoli* grown in plant against *R. phaseoli* Ch24-10 with community grown in plants, using a) up-regulated genes b) down-regulated genes.

## DISCUSSION

The functional synergism that may lead to growth promotion by the tested community in roots may reflect interactions resulting from a long evolutionary history of bacteria which might have been selected as advantageous for plants in roots. Metatranscriptomic analyses seem justified to try to understand the bacterial interactions on roots, and for simplicity we designed an inoculum with few species to be used.

We found that the plants with the multi-species inoculum were statistically larger in comparison with the non-inoculated control and with the plants inoculated with a single organism or with non-inoculated controls. These results suggest that there was synergism between the strains inoculated in corn, which favors the biomass gained by the plant with the multispecies inoculum. In the single plant assays with rhizobia, our results confirm previous experiments with rhizobia in non-legumes that show growth promotion in plants. (43, 44).

Though it is known that various microorganisms favor the growth of crops, less is known on the cooperative interactions among them to produce additional effects on plant growth (2, 5, 15, 45, 46).

From the metatranscriptomic analysis, we found genes encoding transporters showing differential expression under distinct experimental conditions. Transporters belonging to the ATP-binding cassette (or ABC) superfamily couple the energy released from ATP hydrolysis to the translocation of a wide variety of substances into or out of cells (47). We found up regulated genes in *R. phaseoli* in plant encoding ABC transporter for putrescine, which is a polyamine involved in response to osmotic stress. Osmolarity is expected to be high in the rhizoplane. Rapid bacterial adaptation to the increase in osmolarity would facilitate rhizosphere colonization (48)) and allows the adaptation of bacteria to hyperosmolar environments. Accordingly, this transporter has been reported under hyperosmolar conditions in several bacteria such as *Azospirillum* sp. B510, *Burkholderia phytofirmans* PsJN, *Methylobacterium populi* BJ001, *Pseudomonas putida* W619 when they are associated with different plants (49).

Receptor signaling in bacterial chemotaxis occurs within stable complexes between membrane receptors and the CheA kinase, CheB, CheR and CheY kinases. This family of methyl-accepting chemotaxis proteins (MCPs) seem to be important elements in the chemotaxis process (50) When comparing *R. phaseoli* grown in PY medium to *R. phaseoli* Ch24-10 grown in plants after 5 days some down-regulated genes were those involved with chemotaxis, such as those encoding chemotaxis protein CheR, chemotaxis protein CheW. Once on plant roots, rhizobia may no longer need chemotaxis to other sources of nutrients. Additional down-regulated genes were those encoding flagellar synthesis. It is known that molecules of the bacterial envelope such as flagellar proteins, exopolysaccharides, adhesins and lipopolysaccharides are the first structures that contact the host and may be required for bacterial attachment on root surfaces at early times (51).

### Nitrogen fixation and bacterial adaptation to microoxic environment

Nitrogen fixation is a key service in plant ecology but an energetically expensive process. Thus, diazotrophs are normally a minority among bacterial communities in plants (52, 53), and nitrogen fixation is a tightly regulated process which is turned off when fixed nitrogen is available. Therefore, we used a nitrogen free medium to grow maize plants to allow bacterial nitrogen fixing genes to be fully expressed. To promote nitrogen fixation, we included three different diazotrophs in the bacterial maize inoculant.

In previous studies, when *nif* gene expression was analyzed to identify diazotrophs in non-legume roots with a culture independent approach, rhizobial *nif* genes sequences were found to be expressed in rice and sugarcane (54), suggesting that nitrogen fixation by rhizobia may occur out of nodules in non-legume plants (55) As these experiments were performed in field conditions, there were most probably other bacteria that colonized roots in addition to rhizobia (54), but the role of such bacteria on rhizobial *nif* gene expression was not explored. The synthetic ecology approach that we followed here was an attempt to answer this question. Surprisingly, our results showed that *Bacillus* and *Azospirillum* were the front-runners in the interstrain competition for the maize rhizosphere colonization under the conditions tested. Remarkably in rhizobial research, interstrain competition assays for nodule formation are the most valuable tests to define symbiotic defects. However, low levels of nitrogen fixation may occur with *R. phaseoli* Ch24-10 in maize though not detected by acetylene reduction, since Ch24-10 promotes plant growth in the absence of added nitrogen.

*R. phaseoli nif* genes were found expressed in plant when it was inoculated by itself. Furthermore, bacterial respiration genes under low oxygen were found expressed as well. Bacterial respiration is directly related to the supply of ATP; this ATP is necessary to give enough energy for the nitrogen fixation to take place. Nitrogen-fixing bacteria use cbb3-type oxidases encoded by the *fixNOQP* operon and this was expressed as well (56, 57). (58). In spite of rhizobium nitrogen fixation being transcriptionally active no acetylene reduction was detected from these plants associated bacteria. Clearly there are other elements that are needed for *Rhizobium* to fix nitrogen out of the nodule. It was unfortunate that when *Rhizobium* was inoculated with the other bacteria, its nitrogen-fixation machinery was not expressed. Only one of the nitrogen-fixing bacteria from the multispecies inoculum proved functional in plants, namely *Azospirillum*.

Acetylene reduction was detected in *Azospirillum* cultures and when *Azospirillum* was as a single inoculum in plant and more activity was recorded when in combination with *Bacillus*. We surmise that in plant the large *Bacillus* populations attained inhibited *R. phaseoli* Ch24-10, *S. americanum* and *Methylobacterium,* but not *Azospirillum*. However, *in vitro* tests in culture medium this was not the case, since there was no antagonism amongst bacteria. Maybe in the presence of plant roots or in the presence of other bacteria *Bacillus* produces a distinct profile of antibiotics antagonistic to rhizobia and methylobacteria, but further experimentation is needed to test this hypothesis.

It seems plausible that *Bacillus* provides suitable conditions for *Azospirillum* to fix nitrogen, perhaps by inhibiting other bacteria that may compete for nutrients or by making a combined biofilm or by providing low oxygen conditions. This should be further explored. We may even speculate that *Bacillus* may stimulate the excretion of fixed nitrogen by *Azospirillum*. Notably excretion ammonium mutants of nitrogen fixing bacteria are capable of promoting plant growth in nitrogen deficient conditions. Enhanced biofilm formation was observed in multispecies biofilms in marine systems (59), and nitrogen fixation may be stimulated in interspecies biofilms. There are examples of metabolic complementation that may occur between distinct symbiont species in laboratory flow chambers (60). Our results are a basis to further evaluate *Bacillus*/*Azospirillum* inoculum in maize fields to promote nitrogen fixation.

## ACKNOWLEDGEMENTS

This work was supported by Grants from Consejo Nacional de Ciencia y Tecnología (CONACyT) Basic Science grant 253116 and Programa de Apoyo a Proyectos de Investigación e Innovación Tecnológica (PAPIIT) IN207718. We thank Julio Martínez-Romero, for his for technical help. We also thank M. Dunn for critically proofreadingthe manuscript. Lorena Jacqueline Gómez-Godínez is a student from Programa de Doctorado en Ciencias Biomédicas, Universidad Nacional Autónoma de México (UNAM) and was supported by CONACyT (CVU 419877).

## REFERENCES

1. Akhtar N, Arshad I, Shakir MA, Qureshi MA, Sehrish J, Ali L. 2013. Co-inoculation with Rhizobium and Bacillus sp to improve the phosphorus availability and yield of wheat (Triticum aestivum L.). J Anim Plant Sci.

2. Rajendran G, Sing F, Desai AJ, Archana G. 2008. Enhanced growth and nodulation of pigeon pea by co-inoculation of Bacillus strains with Rhizobium spp. Bioresour Technol.

3. Yahalom E, Okon Y, Dovrat A. 1987. Azospirillum effects on susceptibility to Rhizobium nodulation and on nitrogen fixation of several forage legumes. Can J Microbiol 33:510–514.

4. Groppa MD, Zawoznik MS, Tomaro ML. 1998. Effect of co-inoculation with Bradyrhizobium japonicum and Azospirillum brasilense on soybean plants. Eur J Soil Biol.

5. Mansotra P, Sharma P, Sharma S. 2015. Bioaugmentation of Mesorhizobium cicer, Pseudomonas spp. and Piriformospora indica for Sustainable Chickpea Production. Physiol Mol Biol Plants.

6. Tarrand JJ, Krieg NR, Döbereiner J. 1978. A taxonomic study of the Spirillum lipoferum group, with descriptions of a new genus, Azospirillum gen. nov. and two species, Azospirillum lipoferum (Beijerinck) comb. nov. and Azospirillum brasilense sp. nov. Can J Microbiol 24:967–980.

7. Steenhoudt O, Vanderleyden J. 2000. Azospirillum, a free-living nitrogen-fixing bacterium closely associated with grasses: Genetic, biochemical and ecological aspects. FEMS Microbiol Rev.

8. Bashan Y, de-Bashan LE. 2010. How the plant growth-promoting bacterium azospirillum promotes plant growth-a critical assessmentAdvances in Agronomy.

9. Marks BB, Megías M, Ollero FJ, Nogueira MA, Araujo RS, Hungria M. 2015. Maize growth promotion by inoculation with Azospirillum brasilense and metabolites of Rhizobium tropici enriched on lipo-chitooligosaccharides (LCOs). AMB Express.

10. Bano Q, Ilyas N, Bano A, Zafar N, Akram A, Hassan FUL. 2013. Effect of Azospirillum inoculation on maize (Zea mays L.) under drought stress. Pakistan J Bot.

11. Somers E, Ptacek D, Gysegom P, Srinivasan M, Vanderleyden J. 2005. Azospirillum brasilense produces the auxin-like phenylacetic acid by using the key enzyme for indole-3-acetic acid biosynthesis. Appl Environ Microbiol.

12. Cassán F, Vanderleyden J, Spaepen S. 2014. Physiological and Agronomical Aspects of Phytohormone Production by Model Plant-Growth-Promoting Rhizobacteria (PGPR) Belonging to the Genus Azospirillum. J Plant Growth Regul.

13. Dobbelaere S, Croonenborghs A, Thys A, Vande Broek A, Vanderleyden J. 1999. Phytostimulatory effect of Azospirillum brasilense wild type and mutant strains altered in IAA production on wheat. Plant Soil.

14. Idris EE, Iglesias DJ, Talon M, Borriss R. 2007. Tryptophan-Dependent Production of Indole-3-Acetic Acid (IAA) Affects Level of Plant Growth Promotion by Bacillus amyloliquefaciens FZB42. Mol Plant-Microbe Interact.

15. Chen L, Liu Y, Wu G, Veronican Njeri K, Shen Q, Zhang N, Zhang R. 2016. Induced maize salt tolerance by rhizosphere inoculation of Bacillus amyloliquefaciens SQR9. Physiol Plant.

16. Vardharajula S, Zulfikar Ali S, Grover M, Reddy G, Bandi V. 2011. Drought-tolerant plant growth promoting Bacillus spp.: effect on growth, osmolytes, and antioxidant status of maize under drought stress. J Plant Interact.

17. Talboys PJ, Owen DW, Healey JR, Withers PJA, Jones DL. 2014. Auxin secretion by Bacillus amyloliquefaciens FZB42 both stimulates root exudation and limits phosphorus uptake in Triticum aestivium. BMC Plant Biol.

18. López-López A, Rogel MA, Ormeño-Orrillo E, Martínez-Romero J, Martínez-Romero E. 2010. Phaseolus vulgaris seed-borne endophytic community with novel bacterial species such as Rhizobium endophyticum sp. nov. Syst Appl Microbiol.

19. Morgado LB, Willey RW. 2003. Effects of plant population and nitrogen fertilizer on yield and efficiency of maize-bean intercropping. Pesqui Agropecu Bras.

20. Schauer S, Kämpfer P, Wellner S, Spröer C, Kutschera U. 2011. Methylobacterium marchantiae sp. nov., a pinkpigmented, facultatively methylotrophic bacterium isolated from the thallus of a liverwort. Int J Syst Evol Microbiol.

21. Kutschera U. 2007. Plant-associated methylobacteria as co-evolved phytosymbionts: A hypothesis. Plant Signal Behav.

22. Cabanes D, Boistard P, Batut J. 2000. Identification of Sinorhizobium meliloti genes regulated during symbiosis. J Bacteriol.

23. Dent D, Cocking E. 2017. Establishing symbiotic nitrogen fixation in cereals and other non-legume crops: The Greener Nitrogen Revolution. Agric Food Secur.

24. Matsuoka Y, Vigouroux Y, Goodman MM, Sanchez G. J, Buckler E, Doebley J. 2002. A single domestication for maize shown by multilocus microsatellite genotyping. Proc Natl Acad Sci.

25. Xie S, Wu H, Chen L, Zang H, Xie Y, Gao X. 2015. Transcriptome profiling of Bacillus subtilis OKB105 in response to rice seedlings. BMC Microbiol.

26. Camilios-Neto D, Bonato P, Wassem R, Tadra-Sfeir MZ, Brusamarello-Santos LCC, Valdameri G, Donatti L, Faoro H, Weiss VA, Chubatsu LS, Pedrosa FO, Souza EM. 2014. Dual RNA-seq transcriptional analysis of wheat roots colonized by Azospirillum brasilense reveals up-regulation of nutrient acquisition and cell cycle genes. BMC Genomics.

27. Li Y, Tian CF, Chen WF, Wang L, Sui XH, Chen WX. 2013. High-Resolution Transcriptomic Analyses of Sinorhizobium sp. NGR234 Bacteroids in Determinate Nodules of Vigna unguiculata and Indeterminate Nodules of Leucaena leucocephala. PLoS One.

28. Yuan S, Li R, Chen S, Chen H, Zhang C, Chen L, Hao Q, Shan Z, Yang Z, Qiu D, Zhang X, Zhou X. 2016. RNA-Seq Analysis of Differential Gene Expression Responding to Different Rhizobium Strains in Soybean (Glycine max) Roots. Front Plant Sci.

29. Fan B, Carvalhais LC, Becker A, Fedoseyenko D, Von Wirén N, Borriss R. 2012. Transcriptomic profiling of Bacillus amyloliquefaciens FZB42 in response to maize root exudates. BMC Microbiol.

30. Drogue B, Sanguin H, Chamam A, Mozar M, Llauro C, Panaud O, Prigent-Combaret C, Picault N, Wisniewski-DyÃ© F. 2014. Plant root transcriptome profiling reveals a strain-dependent response during Azospirillum-rice cooperation. Front Plant Sci.

31. Araújo WL, Santos DS, Dini-Andreote F, Salgueiro-Londoño JK, Camargo-Neves AA, Andreote FD, Dourado MN. 2015. Genes related to antioxidant metabolism are involved in Methylobacterium mesophilicum-soybean interaction. Antonie van Leeuwenhoek, Int J Gen Mol Microbiol.

32. López-Guerrero MG, Ormeño-Orrillo E, Acosta JL, Mendoza-Vargas A, Rogel MA, Ramírez MA, Rosenblueth M, Martínez-Romero J, Martínez-Romero E. 2012. Rhizobial extrachromosomal replicon variability, stability and expression in natural niches. Plasmid.

33. Rosenblueth M, Martínez-Romero E. 2004. Rhizobium etli maize populations and their competitivenes for roof colonization. Arch Microbiol.

34. Anthony C. 1982. The biochemistry of methylotrophs. Academic press London.

35. Toledo I, Lloret L, Martínez-Romero E. 2003. Sinorhizobium americanus sp. nov., a New Sinorhizobium species nodulating native Acacia spp. in Mexico. Syst Appl Microbiol.

36. Lullien V, Barker DG, de Lajudie P, Huguet T. 1987. Plant gene expression in effective and ineffective root nodules of alfalfa (Medicago sativa). Plant Mol Biol 9:469–478.

37. Andrews S. 2010. FastQC: A quality control tool for high throughput sequence data. Babraham Bioinforma.

38. Bolger AM, Lohse M, Usadel B. 2014. Trimmomatic: A flexible trimmer for Illumina sequence data. Bioinformatics.

39. Langmean B, Salzberg SL. 2012. Bowtie 2. Nat Methods.

40. Liao Y, Smyth GK, Shi W. 2014. FeatureCounts: An efficient general purpose program for assigning sequence reads to genomic features. Bioinformatics.

41. Robinson MD, McCarthy DJ, Smyth GK. 2010. edgeR: a Bioconductor package for differential expression analysis of digital gene expression data. Bioinformatics.

42. Alexa A and Rahnenfuhrer J. 2016. topGO: Enrichment Analysis for Gene Ontology. R Packag version 2260.

43. Chabot R, Antoun H, Kloepper JW, Beauchamp CJ. 1996. Root colonization of maize and lettuce by bioluminescent Rhizobium leguminosarum biovar phaseoli. Appl Environ Microbiol.

44. Gutiérrez-Zamora ML, Martínez-Romero E. 2001. Natural endophytic association between Rhizobium etli and maize (Zea mays L.). J Biotechnol.

45. Alagawadi AR, Gaur AC. 1992. Inoculation of Azospirillum brasilense and phosphate-solubilizing bacteria on yield of sorghum [Sorghum bicolor (L.) Moench] in dry land. Trop Agric (Trinidad Tobago).

46. Cassán F, Perrig D, Sgroy V, Masciarelli O, Penna C, Luna V. 2009. Azospirillum brasilense Az39 and Bradyrhizobium japonicum E109, inoculated singly or in combination, promote seed germination and early seedling growth in corn (Zea mays L.) and soybean (Glycine max L.). Eur J Soil Biol.

47. Davidson AL, Chen J. 2004. ATP-Binding Cassette Transporters in Bacteria. Annu Rev Biochem.

48. Miller KJ, Wood JM. 1996. Osmoadaptation by rhizosphere bacteria. Annu Rev Microbiol.

49. Naveed M, Mitter B, Reichenauer TG, Wieczorek K, Sessitsch A. 2014. Increased drought stress resilience of maize through endophytic colonization by Burkholderia phytofirmans PsJN and Enterobacter sp. FD17. Environ Exp Bot.

50. Grebe TW, Stock J. 1998. Bacterial chemotaxis: the five sensors of a bacterium. Curr Biol.

51. Balsanelli E, Tadra-Sfeir MZ, Faoro H, Pankievicz VC, de Baura VA, Pedrosa FO, de Souza EM, Dixon R, Monteiro RA. 2016. Molecular adaptations of Herbaspirillum seropedicae during colonization of the maize rhizosphere. Environ Microbiol.

52. Hartman K, van der Heijden MGA, Roussely-Provent V, Walser JC, Schlaeppi K. 2017. Deciphering composition and function of the root microbiome of a legume plant. Microbiome.

53. Colin Y, Nicolitch O, Van Nostrand JD, Zhou JZ, Turpault MP, Uroz S. 2017. Taxonomic and functional shifts in the beech rhizosphere microbiome across a natural soil toposequence. Sci Rep.

54. Fischer D, Pfitzner B, Schmid M, Simões-Araújo JL, Reis VM, Pereira W, Ormeño-Orrillo E, Hai B, Hofmann A, Schloter M, Martinez-Romero E, Baldani JI, Hartmann A. 2012. Molecular characterisation of the diazotrophic bacterial community in uninoculated and inoculated field-grown sugarcane (Saccharum sp.). Plant Soil.

55. James EK, Baldani JI. 2012. The role of biological nitrogen fixation by non-legumes in the sustainable production of food and biofuels. Plant Soil.

56. Zufferey R, Preisig O, Hennecke H, Thöny-Meyer L. 1996. Assembly and function of the cytochrome cbb3 oxidase subunits in Bradyrhizobium japonicum. J Biol Chem.

57. Lopez O, Morera C, Miranda-Rios J, Girard L, Romero D, Soberon M. 2001. Regulation of gene expression in response to oxygen in Rhizobium etli: Role of FnrN in fixNOQP expression and in symbiotic nitrogen fixation. J Bacteriol.

58. Talbi C, Sánchez C, Hidalgo-Garcia A, González EM, Arrese-Igor C, Girard L, Bedmar EJ, Delgado MJ. 2012. Enhanced expression of Rhizobium etli cbb3 oxidase improves drought tolerance of common bean symbiotic nitrogen fixation. J Exp Bot.

59. Burmølle M, Webb JS, Rao D, Hansen LH, Sørensen SJ, Kjelleberg S. 2006. Enhanced biofilm formation and increased resistance to antimicrobial agents and bacterial invasion are caused by synergistic interactions in multispecies biofilms. Appl Environ Microbiol.

60. Manzano-marı A. 2011. Serratia symbiotica from the aphid Cinara cedri: a missing link from facultative to obligate insect endosymbiont. PLoS Genet.

